# Healthy adults favor stable left/right hand choices over performance at an unconstrained reach-to-grasp task

**DOI:** 10.1101/2023.10.11.561912

**Authors:** Taewon Kim, Ruiwen Zhou, Samah Gassass, Lei Liu, Benjamin A. Philip

**Author notes:** **Corresponding author:** Benjamin A. Philip, PhD, Washington University School of Medicine in St. Louis, 4444 Forest Park Avenue, MSC-8505-66-1, St. Louis, MO 63108-2212, USA, E-mail address.

## Abstract

Reach-to-grasp actions are fundamental to the daily activities of human life, but few methods exist to assess individuals’ reaching and grasping actions in unconstrained environments. The Block Building Task (BBT) provides an opportunity to directly observe and quantify these actions, including left/right hand choices. Here we sought to investigate the motor and non-motor causes of left/right hand choices, and optimize the design of the BBT, by manipulating motor and non-motor difficulty in the BBT’s unconstrained reach-to-grasp task We hypothesized that greater motor and non-motor (e.g. cognitive/perceptual) difficulty would drive increased usage of the dominant hand. To test this hypothesis, we modulated block size (large vs. small) to influence motor difficulty, and model complexity (10 vs. 5 blocks per model) to influence non-motor difficulty, in healthy adults (n=57). We hypothesized that healthy adults with high non-dominant hand performance in a precision drawing task should be more likely to use their non-dominant hand in the BBT. Our data revealed that increased motor and non-motor difficulty led to lower task performance (slower speed), but participants only increased use of their dominant hand only under the most difficult combination of conditions: in other words, participants allowed their performance to degrade before changing hand choices, even though participants were instructed only to optimize performance. These results demonstrate that hand choices during reach-to grasp actions are more stable than motor performance in healthy right-handed adults, but tasks with multifaceted difficulties can drive individuals to rely more on their dominant hand.

**Statements and Declarations:** Dr. Philip and Washington University in St. Louis have a licensing agreement with PlatformSTL to commercialize the iPad app used in this study.

## Introduction

Our everyday actions often involve the use of one hand over the other, which requires an implicit or explicit decision about which hand to use (Oldfield 1971; Gabbard and Rabb 2000; Sainburg 2002; Scharoun, Scanlan and Bryden 2016). This decision takes on special relevance for patients with unilateral impairment of the upper limb, especially their dominant hand (DH) (Philip, Kaskutas and Mackinnon 2020). Such unilateral impairment occurs frequently after neurological conditions such as stroke (Mani et al. 2013), but also impacts the life of patients with chronic peripheral injuries, e.g. upper extremity peripheral nerve injury (Wojtkiewicz et al. 2015). After peripheral nerve injury to the DH, patients continue using their affected hand whenever possible, despite their uninjured hand being more dexterous (Philip et al. 2022a). Therefore, encouraging compensation with the use of a healthy non-dominant hand (NDH) is crucial for these patients to regain their ability to function normally during daily activities.

Despite the importance of hand dominance and choice for everyday life and rehabilitation, few studies have investigated the factors that influence hand choice during reach-to-grasp actions in healthy adults. Previous work has demonstrated that healthy adults use their NDH to reach to more of the workspace when visual feedback is unavailable (Przybyla et al. 2013); and use their DH to reach to more of the workspace when the reached-to object will be used in a more complex task (e.g. tool use instead of simple pickup) regardless of grasp difficulty (Mamolo et al. 2004; Leconte and Fagard 2006; Bryden, Mayer and Roy 2011; Stone, Bryant and Gonzalez 2013a), or under greater cognitive load (Liang, Wilkinson and Sainburg 2018b). However, it remains unclear how well these results integrate, given their different task contexts. More importantly, none of these studies modified task difficulty in both sensorimotor and cognitive-perceptual aspects, thus leaving unanswered questions about how motor and non-motor difficulties might interact.

The Block Building Task (BBT) provides a means to measure hand choices over time in unconstrained situations, with the potential to modify both motor and non-motor aspects of the task (Gonzalez and Goodale 2009; Stone, Bryant and Gonzalez 2013b). The BBT requires participants to pick up Lego blocks (The Lego Group, Billund, Denmark) and incorporate them into a simple model. The task naturally induces the participant to choose one hand for reaching and grasping each block, followed by bimanual interaction to construct the model. Importantly, this task allows direct quantification of left/right-hand choices for reach-to-grasp action in an unconstrained environment, in the context of complex object manipulations (building the model with the grasped blocks). Recently, this task has been used in peripheral nerve injury patients to illustrate the stability of hand choice after peripheral nerve injuries to the dominant hand, and it is sensitive to the interaction between peripheral nerve injury and hand dominance (Philip et al. 2022a). Although the BBT has been used as a rapid and feasible measurement tool of hand choice during reaching and grasping actions in unconstrained environments, few studies have yet investigated what aspects of the task current version of BBT is not designed to modulate hand choices in reach-to-grasp action, therefore, it is unclear what aspects of the task drive its specific results.

In this study, we aimed to investigate how motor and non-motor difficulty factors drive left/right hand choices in an unconstrained reach-to-grasp context, and specifically how motor and non-motor factors can modulate task difficulty in the BBT. We hypothesized that greater motor and non-motor (e.g. cognitive/perceptual) difficulty would drive increased usage of the DH during reaching and grasping. We modulated motor difficulty in the BBT by changing block sizes (Small vs. Large), and non-motor difficulty by changing model complexity (10 vs. 5 blocks per model). In addition, we hypothesized that NDH use would be correlated with NDH performance in precision drawing task (Philip et al. 2023).

## Methods

### Study overview

This was cross-sectional single-arm study involving a single laboratory visit. The study was approved by the local Institutional Review Board ethics committee and all participants gave informed consent before participating in the study. Data were stored and managed via the Research Electronic Data Capture system (Harris et al. 2009).

### Participants

Fifty-seven right-handed healthy adults (3 males), aged between 22 and 51 (28 ± 7 years), participated in this experiment. Participants were recruited without balancing genders because previous studies revealed no gender difference in the Block Building Task (BBT) (Gonzalez and Goodale 2009). The inclusion criteria were: right handed as determined by Edinburgh Handedness Inventory score 50+ (Oldfield 1971), normal or corrected-to-normal vision (self-report) and fluency in speaking, reading, and writing English. Exclusion criteria included individuals who have motor disabilities or impairments affecting the upper limb (self-report), and those who would be unable to come to campus to complete the tasks on site. All participants were recruited from the Washington University in St. Louis community to minimize risks during the COVID-19 pandemic (i.e. recruited from individuals who were already coming onto campus).

Sample size was determined from a power analysis based on preliminary data from a subset of the participants (n=19). Based on the mean and variance-covariance matrices in the preliminary data, for each precision drawing variable (see below) to distinguish between the 4 BBT variants, we needed 33-40 participants to achieve power ≥ 0.8. Based on this power analysis, participants were recruited toward an n=40 goal with an additional 10% margin in case of screening failures, leading to n=44.

Thirteen additional participants had previously been recruited to perform an identical version of the BBT without the precision drawing task. These participants were included in our primary analyses (i.e. all analyses that do not mention the precision drawing task), for a total n=57.

### Materials and procedures

#### Block building task

Participants were seated in front of a table with 40 Lego blocks and instructed to build different “models” one at a time until all 40 blocks were used, as shown in **Fig. 1**. The models were presented in a counterbalanced order. Importantly, the BBT was repeated 4 times for each participant, each repetition using a different variant to modulate task difficulty. The four variants were defined by a 2x2 design that varied motor difficulty via Block Size (Normal, 2.2 ± 1.4 cm3; vs. Small, 0.8 ± 1.6 cm3), and non-motor difficulty via Quantity (10 vs. 5 blocks per model), with examples shown in **Fig. 2**. Variants were presented in counterbalanced order. Block locations on the table were different for each variant, consistent across participants. Participants were instructed to build the model as quickly and accurately as possible; participants received no cues about how to use their hands. Performance was video recorded.

**Figure 1:**
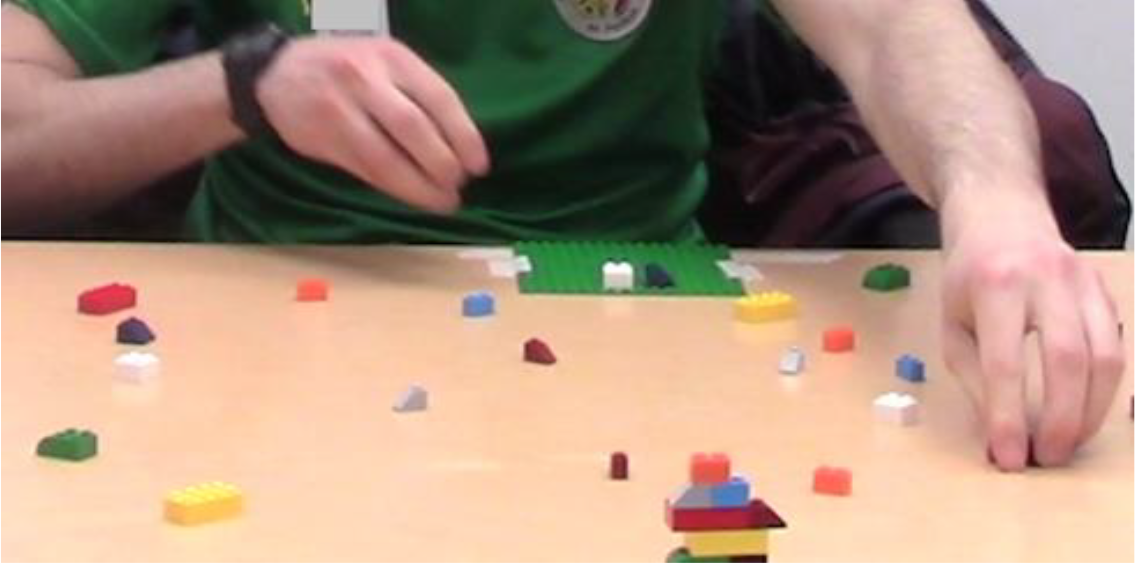
Block Building Task. Figure reproduced from Philip et al (2022). Participants build 4 models (bottom center) from 40 Lego bricks. Participants are instructed to build quickly and accurately, but receive no instructions about hand use.

**Figure 2.**
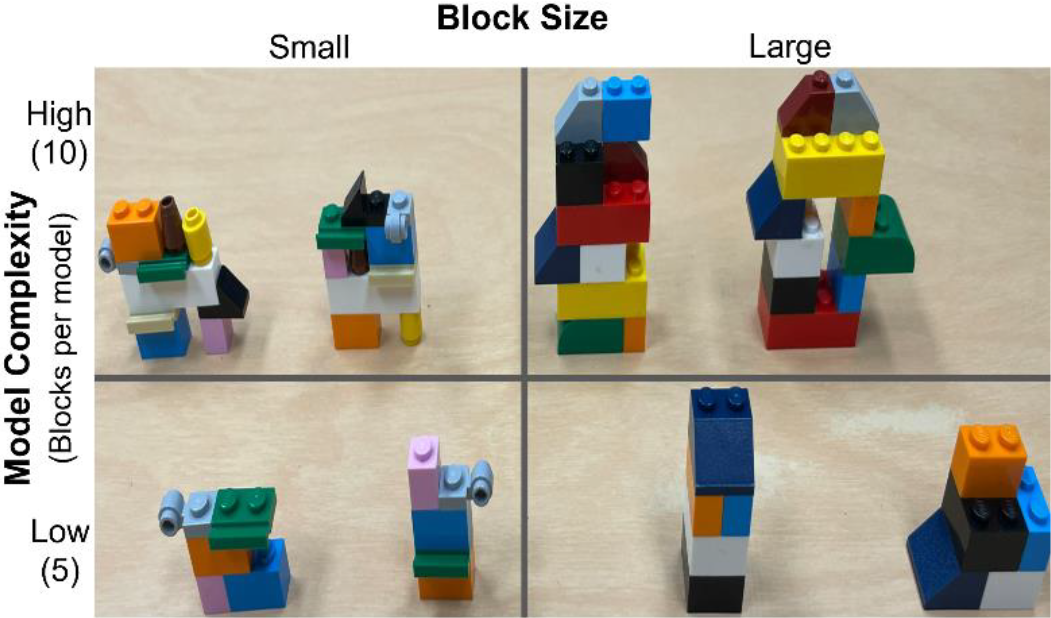
Example models, illustrating 2 × 2 design of motor and non-motor difficulty. Motor difficulty was modulated via Block Size (Large vs. Small), and non-motor difficulty Model Complexity (10 vs 5 blocks per model).

Outcome measures were: fraction of grasps with both hand and speed (blocks/sec). To assess these, each video was reviewed by two raters using BORIS event logging software (Friard and Gamba 2016). In cases where the two raters disagreed on any of the participant’s left/right choices, or their measurements of any model’s start/end time differed by >1.0 sec, the two raters reviewed the video together to reach a consensus. Otherwise, start/end times were averaged between the two raters for each model, and then divided by the number of blocks per model to determine the speed. During study preparation, preliminary tests were performed on variants that involved different kinds of gloves to modify tactile feedback. In small preliminary samples (n=5-15), these variants did not produce consistent trends toward effects on hand choice; as a result, those variants were not tested or analyzed further.

#### Precision drawing task (motor performance)

After completing the BBT, participants performed a precision drawing task using both hands, starting with the right dominant hand. Precision drawing performance was measured via the iPad STEGA app (PlatformsSTL, St. Louis, MO, USA). This app has been successfully employed to measure a precision drawing skill (Philip et al. 2023), and is based on a precision drawing task with a history of motor neuroscience research in healthy and clinical populations (Philip and Frey 2014; Philip and Frey 2016; Philip, McAvoy and Frey 2021). In the STEGA app, participants used an iPad 6th Generation and Apple Pencil (Apple Inc, Cupertino, CA, USA) to draw within the bounds of abstract symmetrical shapes. Each participant completed a total of 30 trials with each hand, comprising 15 shapes at two levels of difficulty each (5mm or 6mm tolerance). Participants were instructed to complete each shape as quickly as possible while staying within the bounds.

The STEGA app collected raw data at a sampling rate of 50Hz on pen position (0.5 mm precision), position and angular errors (deviation 156 from ideal line, measured in mm and degrees respectively), and time (measured in milliseconds). From these raw data, four primary dependent variables were derived: (1) speed (mm/s, mean of each trial), (2, 3) position accuracy (-1 * position error), (3) direction accuracy (-1 * angular error), and (4) velocity smoothness, quantified as a number of submovements per unit distance (-1 * velocity peaks per shape part) to capture motor performance. These four dependent variables were chosen because they represent movement characteristics that are specialized to either hand (Mutha, Haaland and Sainburg 2012).

#### Data Analysis

Data analyses were performed in R version 4.2.1 (R Foundation, Vienna, Austria). To identify which variants, motor and non-motor task difficulty can modulate hand choice and speed during reach-to-grasp action, we performed 2 (Block Size: Normal, Small) by 2 (Model Complexity: 10, 5 blocks per model) repeated measure ANOVA, separately for each outcome variable. In addition, we carried out a correlation coefficient analysis between motor and non-motor variants and behavioral variables including velocity, direction, and distance in drawing performance to assess correlation with hand choice in BBT for any measure of drawing performance. Statistical significance was detected at a=0.05.

## Results

### Dominant hand use increased only under the combination of small blocks and high model complexity

We measured the influences on hand usage (fraction of grasps performed with the DH) during the BBT (block building task) with a 2 (Block Size: Small vs Large) x 2 (Model Complexity: 10 vs 5 blocks per model) repeated measured ANOVA. We found that hand usage depended on the interaction between Block Size and Model Complexity, as shown in **Fig. 3A**. Specifically, for the dependent variable of DH usage, we found no significant main effect of Block Size, F (1, 56) = 3.54, *P* = 0.0617, or Model Complexity, F (1, 56) = 0.11, *P* = 0.7402. However, we found significant Block Size × Model complexity interaction, F (1, 56) = 10.93, *P* = 0.0012. Post-hoc analyses revealed that the interaction effect arose because Block Size had a significant effect on hand usage during high Model Complexity (Block Size (p < 0.05, estimated coefficient - 0.0471), t (54) = -4.429, but not during low Model Complexity (p = 0.327, coefficient 0.0129), t (54) = 0.99. These findings indicate that block size did not matter at low model complexity (5 blocks); but at high model complexity (10 blocks), small blocks led to significantly higher DH usage. In other words, participants used the DH more in the highest-difficulty condition (high model complexity and small blocks) compared to all other conditions.

**Figure 3.**
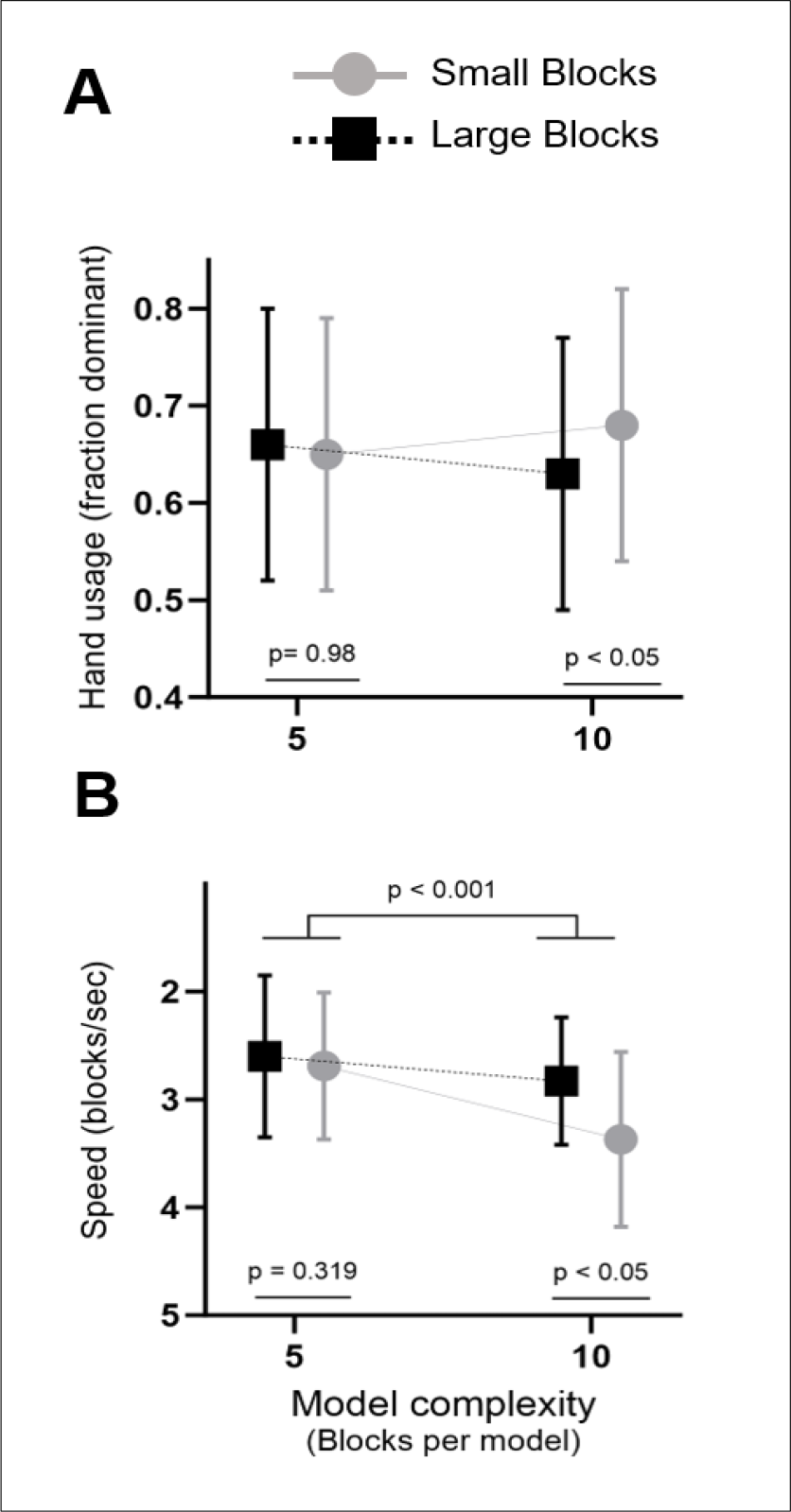
Effects of Block Size and Model Complexity on hand usage and speed. A: Hand usage had no main effects, but an interaction effect: DH usage increased only under the combination of small blocks and high complexity. B: Speed had main effects of Block Size and Model Complexity, with an interaction effect such that small Block Size only decreased speed under high Model Complexity.

### Speed increased only under the high model complexity in both block sizes

We measured the influences on speed (seconds per block) during the BBT with another 2 × 2 repeated measures ANOVA, identical to the above except for the different outcome variable (speed). We found that speed depended on both Block Size, Model Complexity, and their interaction, as shown in **Fig. 3B**. Specifically, we found significant main effects for Block Size, F (1, 56) = 20.42, *P* = 1.19e-05; Model complexity, F (1, 56) = 44.09, *P* = 4.52e-07; and interaction, F (1, 56) = 10.81, *P* = 0.00124. We performed two post-hoc analyses on the interaction effect. Our post-hoc examination of the Block Size effect revealed that the small block size was associated with lower speed during high Model Complexity (p < 0.05, estimated coefficient of -0.5347, t (54) = -5.48548), but Block Size did not affect speed during low Model Complexity (p = 0.3861, estimated coefficient -0.0843, t (54) = -0.873812). Our post-hoc examination of the Model Complexity effect revealed that it had a significant effect on speed during both Block Sizes, small (p < 0.05, estimated coefficient -0.68, t (54) = -7.804) and large (p < 0.05, estimated coefficient -0.2296, t (54) = - 2.037. Therefore, increased model complexity always induced lower speed, and this effect was magnified for small blocks.

### Block building task performance did not correlate with precision drawing performance

To investigate the relationship between hand usage and hand function, participants completed a precision drawing task with each hand. We examined the correlations between the four BBT variants (2 Block Sizes x 2 Model Complexities) and four behavioral variables from the STEGA precision drawing task (speed, position accuracy, direction accuracy, and velocity smoothness). Contrary to our initial hypothesis, we found no significant correlation between any measure of drawing performance and any BBT variants (|r| < 0.4, p > 0.05), as shown in **Fig. 4**.

**Figure 4.**
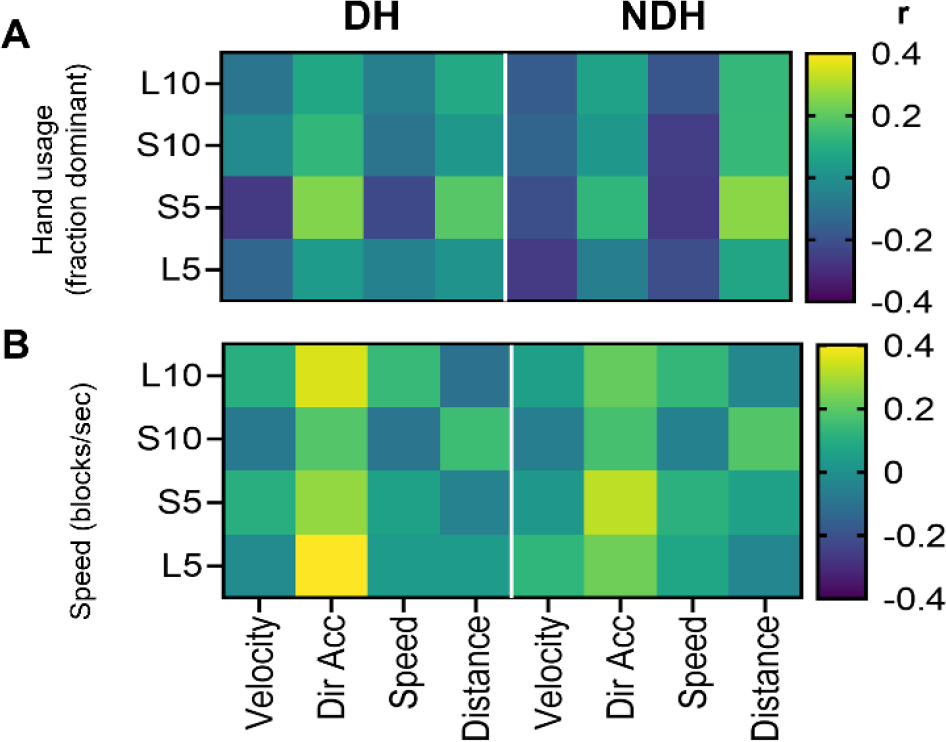
No significant correlations between precision drawing task and BBT. For each hand (DH and NDH), and all four BBT conditions (rows), none of the four precision drawing variables correlated significantly with (A) hand usage or (B) speed.

### Covariance between hand usage and speed did not explain Block Building Task performance

To determine whether our above results were influenced by possible covariance between our outcome measures (hand usage, speed), we performed a multivariate analysis on the two outcome measures together. Our multivariate linear mixed model identified significant effects of block size (estimated coefficient -0.056, p < 0.001), model complexity, (estimated coefficient -0.038, p = 0.002) and their interaction (estimated coefficient 0.067, p < 0.001). Therefore, both factors still had significant effects on performance, even when analyzing both outcome measures together. Moreover, the two outcome measures were only weakly correlated with each other, r ≤ 0.37 in all variants, as shown in **Fig. 5**. Therefore, our results are unlikely to be an artifact of covariance or other interdependence between our outcome measures.

**Figure 5.**
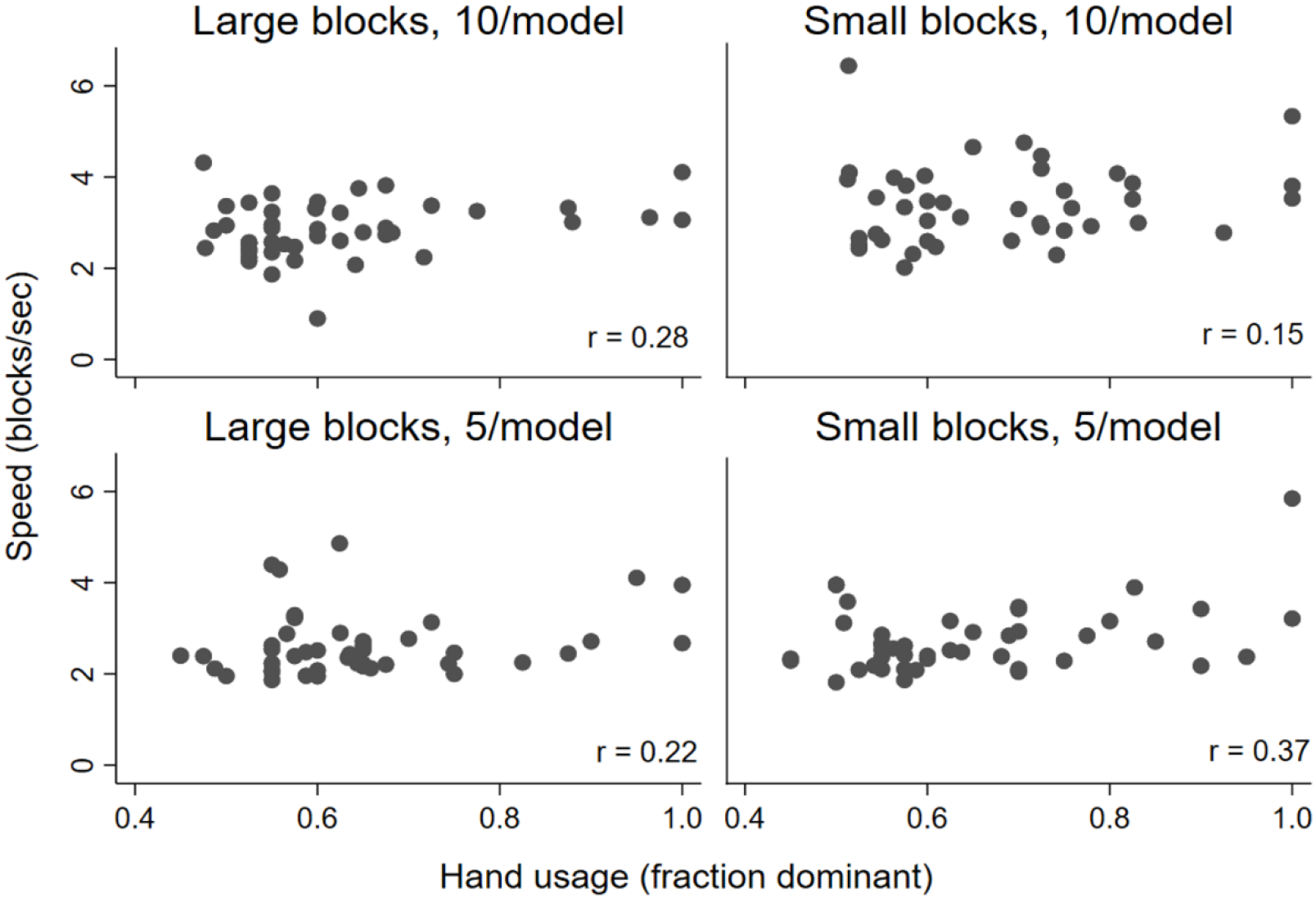
Hand usage and speed were only weakly correlated (r ≤ 0.37), for all variants of the BBT.

## Discussion

The present study represents one of the first systematic attempts to modulate hand usage by controlling the difficulty of an unconstrained reach-to-grasp task. We created and tested new variants of the block building task (BBT) to identify features that could serve as task difficulty axes to modulate hand usage for reach-to-grasp action, and/or hand usage’s relationship with motor performance. We found that participant performance (e.g. speed) responded to both block size (motor difficulty) and model complexity (non-motor difficulty), but hand usage (left/right hand choices) only changed when participants experienced the most difficult combination of these factors – even though participants were instructed to prioritize speed and accuracy, not hand choices. This illustrates the separability of hand performance and choice, and that hand choice preferences in healthy adults are more stable than other aspects of motor behavior.

### Increased cognitive-perceptual difficulty led to slower performance, but only influenced hand usage under high motor difficulty

In the present study, we controlled model complexity to manipulate non-motor difficulty, which we classify as “cognitive-perceptual” difficulty. Greater model complexity should require higher levels of visual-spatial processing and working memory to support the (otherwise unchanged) elaborate fine motor skills of building or assembling blocks. We expected cognitive-perceptual difficulty to influence hand choice because higher-demand cognitive tasks require greater recruitment of cognitive resources such as attention, perception and working memory (Sauseng et al. 2010), and individuals select their dominant hand more frequently as cognitive load increases (Liang, Wilkinson and Sainburg 2018b). This phenomenon is thought to reflect cognitive visual-spatial processing in movement, where more complex task demands are associated with greater information processing demands (Rosenbaum 1980; Stelmach 2014), which lead to longer reaction time (Bootsma et al. 2018) and increased use of the dominant hand (Bryden, Mayer and Roy 2011; Liang, Wilkinson and Sainburg 2018a; Liang, Wilkinson and Sainburg 2018b). However we found here that increased cognitive-perceptual difficulty in the BBT did not directly affect hand choice. Instead, increased cognitive-perceptual difficulty only led to increased DH use under conditions of high grasp motor difficulty.

Negative results are sometimes difficult to interpret, but our multiple task variants and outcome variables allow us to rule out a number of alternatives. Conceivably, our cognitive-perceptual manipulation could have been insufficient to affect participant behavior; however, our cognitive-perceptual manipulation (model complexity) always affected movement speed. Alternatively, it is conceivable that our negative result arose as an artifact of statistical thresholds; however, this interpretation is unlikely given the different direction of the model-complexity effect at each block size (compare two slopes in **Fig. 3A**). One uncertainty cannot be resolved by our current data: the interaction between motor and non-motor difficulty may be specific to the combination of those two factors or may represent a simpler “total difficulty” effect.

Regardless of the source of our difficulty effect, it had a greater impact on task performance (speed) rather than usage, despite instructions to the contrary: participants were instructed to move quickly and accurately, but their instructions avoided any mention of hand choice. (“Accuracy” at the BBT effectively also means speed: participants continued the task until they completed all movements, so movement errors would introduce delays as the participant made additional corrective actions.) This extends previous findings about the stability of hand choice: we have previously shown that people avoid *decreasing* DH use even when it would be useful to do so (Philip et al. 2022b), but here we demonstrate that people also avoid *increasing* DH use.

### Motor difficulty effects could depend on grasping itself or the post-grasping task

We found that motor difficulty (block size) affected hand choice and movement speed during conditions of high model complexity (10 blocks per model). This interaction effect has been discussed in the previous section, but it is important to briefly discuss the effects of block size on their own. Previous studies had found that DH use was increased by more difficult post-grasp actions, but not by more difficult grasp itself (Mamolo et al. 2004; Leconte and Fagard 2006; Bryden, Mayer and Roy 2011). A limitation of the BBT is that it is difficult to separate these two types of motor difficulty: because the BBT presents a context in which reach-to-grasp actions occur as part of an unconstrained goal-directed task, smaller blocks have two effects: they increase difficulty of the initial grasp and also the subsequent model-building. Future studies could disambiguate this by quantifying hand usage during the model-building phase of the BBT, where we would expect a stronger effect of block size. Given the prior research, it seems likely that the current data’s motor difficulty interaction effect arose primarily from difficulty in the post-grasp task (i.e. model building).

### Hand choices did not correlate with precision drawing performance

We found no relationships between hand choices and precision drawing performance, for any version of the BBT or any precision drawing variable. In other words, our data did not support our hypothesis that participants with better NDH control would be more willing to use their NDH. This could potentially because the two tasks were not well aligned: reach-to-grasp performance may be unrelated to precision endpoint (drawing) control performance. Indeed, these two tasks have separable performance characteristics (Pacilli et al. 2014; Israely and Carmeli 2017). For example, even if individuals with better NDH drawing performance are more likely to use that hand for fine manipulation, this might not affect their reach-to-grasp choices. Future studies should pair the BBT with other tests of manual dexterity to assess other aspects of motor performance. With our current measures, our precision drawing data support the idea that left/right hand choices for reaching do not depend on fine endpoint control capacity among healthy adults. This is consistent with our other data showing that hand usage is poorly correlated with task performance even within the same reach-to-grasp task.

## Conclusion

In this study, we used Block Building Task to identify the effects of motor and non-motor difficulty on hand usage for reach-to-grasp actions. We found that hand usage (left/right hand choices) was relatively insensitive to task difficulty: participants increased the use of their dominant hand only for the most difficult combination of conditions, even though both motor and non-motor difficulty influenced task performance (speed). Hand usage was independent from performance of the reach-to-grasp task, or performance at a precision drawing task.

These results demonstrate the stability of left/right hand choices in healthy adults, even against increased use of the dominant hand. Participants were told to prioritize performance speed, but nevertheless sacrificed speed in favor of maintaining a consistent ratio of left/right hand choices across the workspace.

## Acknowledgements

This work was funded by NIH/NINDS R01 NS114046 to BAP. The authors would like to thank Setsu Uzume for their help with data collection; and Xinye Wu, Jackie Hardy, Setsu Uzume, Delaney McIntyre, Tina Nguyen, Rachel Graves, and Elizabeth Magee for their help with video coding.

## Data availability

The data from the current study are available from the corresponding author on request.

## References

Bootsma JM, Hortobágyi T, Rothwell JC, Caljouw SR (2018) The role of task difficulty in learning a visuomotor skill. Medicine and Science in Sports and Exercise 50:1842–1849

Bryden PJ, Mayer M, Roy EA (2011) Influences of task complexity, object location, and object type on hand selection in reaching in left and right-handed children and adults. Developmental Psychobiology 53:47–58 doi: 10.1002/dev.20486

Friard O, Gamba M (2016) BORIS: a free, versatile open‐source event‐logging software for video/audio coding and live observations. Methods in Ecology and Evolution 7:1325–1330

Gabbard C, Rabb C (2000) What determines choice of limb for unimanual reaching movements? The Journal of General Psychology 127:178–184

Gonzalez CL, Goodale MA (2009) Hand preference for precision grasping predicts language lateralization. Neuropsychologia 47:3182–3189

Harris PA, Taylor R, Thielke R, Payne J, Gonzalez N, Conde JG (2009) Research electronic data capture (REDCap)—a metadata-driven methodology and workflow process for providing translational research informatics support. Journal of biomedical informatics 42:377–381 doi: 10.1016/j.jbi.2008.08.010

Israely S, Carmeli E (2017) Handwriting performance versus arm forward reach and grasp abilities among poststroke patients, a case-control study. Top Stroke Rehabil 24:5–11 doi: 10.1080/10749357.2016.1183383

Leconte P, Fagard J (2006) Which factors affect hand selection in children’s grasping in hemispace? Combined effects of task demand and motor dominance. Brain Cogn 60:88–93 doi: 10.1016/j.bandc.2005.09.009

Liang J, Wilkinson K, Sainburg RL (2018a) Is Hand selection modulated by cognitive–perceptual load? Neuroscience 369:363–373

Liang J, Wilkinson KM, Sainburg RL (2018b) Cognitive-perceptual load modulates hand selection in lefthanders to a greater extent than in right-handers. Exp Brain Res 237:389–399 doi: 10.1007/s00221-018-5423-z

Mamolo CM, Roy EA, Bryden PJ, Rohr LE (2004) The effects of skill demands and object position on the distribution of preferred hand reaches. Brain Cogn 55:349–351 doi: 10.1016/j.bandc.2004.02.041

Mani S, Mutha PK, Przybyla A, Haaland KY, Good DC, Sainburg RL (2013) Contralesional motor deficits after unilateral stroke reflect hemisphere-specific control mechanisms. Brain 136:1288–1303

Mutha PK, Haaland KY, Sainburg RL (2012) The effects of brain lateralization on motor control and adaptation. Journal of motor behavior 44:455–469

Oldfield RC (1971) The assessment and analysis of handedness: the Edinburgh inventory. Neuropsychologia 9:97–113

Pacilli A, Germanotta M, Rossi S, Cappa P (2014) Quantification of age-related differences in reaching and circle-drawing using a robotic rehabilitation device. Applied Bionics and Biomechanics 11:91–104

Philip BA, Frey SH (2014) Compensatory changes accompanying chronic forced use of the nondominant hand by unilateral amputees. Journal of Neuroscience 34:3622–3631

Philip BA, Frey SH (2016) Increased functional connectivity between cortical hand areas and praxis network associated with training-related improvements in non-dominant hand precision drawing. Neuropsychologia 87:157–168

Philip BA, Kaskutas V, Mackinnon SE (2020) Impact of handedness on disability after unilateral upper-extremity peripheral nerve disorder. Hand 15:327–334

Philip BA, Li F, Hawkins-Chernof E, Chen L, Swamidass V, Zwir I (2023) Motor Assessment With the STEGA iPad App to Measure Handwriting in Children. The American Journal of Occupational Therapy 77:7703205010

Philip BA, McAvoy MP, Frey SH (2021) Interhemispheric parietal-frontal connectivity predicts the ability to acquire a nondominant hand skill. Brain connectivity 11:308–318

Philip BA, Thompson MR, Baune NA, Hyde M, Mackinnon SE (2022a) Failure to Compensate: Patients With Nerve Injury Use Their Injured Dominant Hand, Even When Their Nondominant Is More Dexterous. Archives of Physical Medicine and Rehabilitation 103:899–907

Philip BA, Thompson MR, Baune NA, Hyde M, Mackinnon SE (2022b) Failure to Compensate: Patients With Nerve Injury Use Their Injured Dominant Hand, Even When Their Nondominant Is More Dexterous. Arch Phys Med Rehabil 103:899–907 doi: 10.1016/j.apmr.2021.10.010

Przybyla A, Coelho CJ, Akpinar S, Kirazci S, Sainburg RL (2013) Sensorimotor performance asymmetries predict hand selection. Neuroscience 228:349–360 doi: 10.1016/j.neuroscience.2012.10.046

Rosenbaum DA (1980) Human movement initiation: specification of arm, direction, and extent. Journal of Experimental Psychology: General 109:444

Sainburg RL (2002) Evidence for a dynamic-dominance hypothesis of handedness. Experimental brain research 142:241–258

Sauseng P, Griesmayr B, Freunberger R, Klimesch W (2010) Control mechanisms in working memory: a possible function of EEG theta oscillations. Neuroscience & Biobehavioral Reviews 34:1015–1022

Scharoun SM, Scanlan KA, Bryden PJ (2016) Hand and grasp selection in a preferential reaching task: the effects of object location, orientation, and task intention. Frontiers in psychology 7:360

Stelmach GE (2014) Information processing in motor control and learning. Academic Press Stone KD, Bryant DC, Gonzalez CL (2013a) Hand use for grasping in a bimanual task: evidence for different roles? Exp Brain Res 224:455–467 doi: 10.1007/s00221-012-3325-z

Stone KD, Bryant DC, Gonzalez CL (2013b) Hand use for grasping in a bimanual task: evidence for different roles? Experimental Brain Research 224:455–467

Wojtkiewicz DM, Saunders J, Domeshek L, Novak CB, Kaskutas V, Mackinnon SE (2015) Social impact of peripheral nerve injuries. Hand 10:161–167

